# Response surface methodology for melanin nanoparticle production optimization from producer strain *Pseudomonas stutzeri* BTCZ305 with invitro anti-inflammatory and wound healing potential

**DOI:** 10.64898/2026.07.08.737209

**Authors:** Dayana Mathew, Sarita. G. Bhat

## Abstract

Culture conditions were optimized for the production of melanin nanoparticle by the bacterial strain *Pseudomonas stutzeri* BTCZ 305. Response surface methodology was employed for determining the most significant fermentation conditions using variables including, pH, temperature and L-tyrosine concentration identified through one-factor-at-a time approach. Box-behnken design consisting of 17 different combinations of all these factors were performed. Using this methodology, a quadratic regression model was built and the optimal combinations of media constituents for maximum melanin production 1192.27 μg/mL were determined as temperature (32.5 °C), pH (8.5) and L-tyrosine concentration (7 g/L). Melanin productionwas obtained experimentally coincident with the predicted value and the model was proven to be adequate. The nanostructural distribution, its stability in colloidal suspension and particle size were also characterized with the help of TEM, particle size analysis and Zeta potential. The potent applicability of this molecule in anti-inflammation and wound healing was also elucidated.

**Highlights:**

- Melanin nanoparticle synthesis by marine bacterial strain *Pseudomonas stutzeri* BTCZ 305
- Media optimization using one-factor-at-a-time method
- Statistical media optimization using Response surface methodology and Box-behnken design of experiment
- Nanoparticle characterization using TEM, particle size analysis and zeta potential. Its functional role in anti-inflammation and wound healing.

## 1. Introduction

Melanins are biopolymers ubiquitously distributed in nature, have shown to be promising among various research areas, like biomedicine, dermo-cosmetics, nanotechnology, and bioengineering. Due to the increased biodegradability and biocompatibility of melanin particles, assume a strong relevance in the production of biomaterials. With the increase in demand for this biopolymer, large-scale fermentation of mesophilic bacteria is gaining importance. Optimization of media by classical one-factor-at-a-time method involves changing one variable at a time while keeping other factors constant. The time-consuming conventional methods for multifactor experimental design like Placket Burman method are incapable of detecting the true optimum, due to the interactions among the factors. On the other hand, response surface methodology (RSM) involves a collection of statistical techniques for designing experiments, building models, evaluating the effects of factors, and searching optimum conditions of factors for desirable responses(Wang *et al*.,2010; Surwase *et al*.,2013; Raman *et al*.,2015 Manivasagan *et al*.,2013; El-Batal *et al*., 2017; El-Naggar *et al*.,2017). This method has been successfully applied in this study for optimization of the melanin nanoparticle production, particularly by *Pseudomonas stutzeri* strain BTCZ305.The application of nano sized melanin particles in molecular imaging, wound healing, drug delivery, photo thermal therapy etc, have been explained in several other studies (Fan *et al*., 2014; Zhou *et al*.,2019; Ju *et al*.,2016)

Wound healing is a vital process, during which specialized immune cells invade the wound site in a specific sequence and seal the lesion to prevent the loss of fluids and infection by foreign organisms and to aid in the regeneration of the tissue (Moeini *et al*.,2020). Here in this study melanin nano particle obtained from marine bacteria *Pseudomonas stutzeri* BTCZ 305 was critically studied with the help of different characterization methods involving UV-Visible spectroscopy, FT-IR, TEM, PSA *etc*. and its vital role in anti-inflammation and wound healing. From the study it was observed that Nano melanin has strong anti-inflammatory activity by inhibiting protein denaturation with IC_50_ value of 52.39 and wound healing potential was estimated using L6 cell line.

## 2. Materials and method

### 2.1 Microorganism and culture conditions

#### 2.1.1 Microorganism

*Pseudomonas stutzeri* strain BTCZ305 isolated during Sagar Sampada cruise no.305 from Arabian sea sediments (9 6’N, 75 22’ E) at 96 meter depth during a previous study was stored in the lab (Kurian *et al*.,2014) using paraffin over lay method was purified and further screened for extracellular melanin production in L-tyrosine agar plates. The promising strains were sorted and used for melanin production in media.

#### 2.1.2 Chemicals used

L-tyrosine, synthetic DOPA melanin from Sigma, USA, sodium chloride (NaCl), magnesium sulfate (MgSO_4_), potassium dihydrogen phosphate (KH_2_PO_4_), and tryptone soy broth (TSB) were obtained from HiMedia Chemicals, India. ethidium bromide, were purchased from Sigma, USA.

### 2.2 Biochemical and molecular identification of bacteria

#### 2.2.1 Biochemical characterization of bacterial strain

Biochemical characterization including Gram staining, Indole, Methyl red, Voges-Proskauer, citrate, catalase, oxidase as well as Sugar utilization using glucose, adonitol, arabinose, lactose sorbitol, mannitol, Rhamnose and sucrose were performed.

#### 2.2.2 Molecular identification and phylogenetic tree construction of the bacterial strain

Genomic DNA was isolated and purified; a portion of the 16S rDNA (1.5kb size) was amplified using a primer pair for 16S rDNA in thermal cycler (BioRad, CA, USA) with universal primers for 16S rDNA [30]. The product after PCR amplification were purified by gene clean kit and the nucleotide sequence was determined by 3730XL DNA ANALYZER using big dye terminator kit and sample files were analysed using Applied Biosystems Sequencing Analysis Software v5.2. The identity of the sequences was determined by comparing the 16S rDNA sequences available in the public nucleotide databases at the National Centre for Biotechnology Information (NCBI) by using their World Wide Web site (http://www.ncbi.nlm.nih.gov) and the BLAST(Basic Local Alignment Search Tool) [1] Algorithm and the nucleotide sequence was submitted to NCBI Genbank. Phylogenetic tree was constructed using Mega X.

### 2.3 Time course study on melanin production

The cell biomass and melanin production kinetics were recorded in batch culture at regular time intervals of 9 days starting from log phase to stationary phase under submerged culture condition was designed. The cell biomass was estimated at regular interval of 12 hrs at OD_600_ nm followed by decellularizing to estimate melanin production. The cell supernatant was removed by centrifugation at 10,000 rpm at 4 °C for 15 min. The resulting supernatant was spectrophotometrically analysed for melanin production at OD_400_ nm.

### 2.4 Selection of most influencing factors in fermentation media through OFAT method and statistical analysis

One-factor-at-a-time method was carried out for determining the most influencing media factors from Yabuuchi and Ohyama’s tyrosine basal broth with 2g/L L-tyrosine (Yabuuchi and Ohyama 1972). (Composition of the media: sodium chloride (NaCl) 5g/L, magnesium sulfate (MgSO_4._ 5H_2_O) 0.1g/L, potassium dihydrogen phosphate (KH_2_PO_4_.7H_2_O). 2g/L at a pH of 8, incubated at 37 °C for 9 days in an environmental orbital shaker (140 rpm). Various culture conditions like initial medium pH (6-10), incubation temperature (30-50 °C) and L-tyrosine concentration (0.2-20 g/L) along with other components in its normal level in media were optimized for enhanced production of melanin by *Pseudomonas stutzeri* BTCZ305 in submerged fermentation condition.

#### 2.4.1 Statistical analysis

Statistically analysis was performed using Graph pad prism 8 in Windows 10. The results are means of three replicates +/-SD.

### 2.5 Statistical media optimization through Response surface methodology and Box-behnken design

#### 2.5.1 Software package

Design of experiment was done by using Design Expert software version13 (Stat-Ease, Minneapolis, MN, USA) and Individual influence of each factor was identified and Optimum culture condition was evaluated and Performance at the optimum condition as well as the interaction effect of factors at the optimum level was estimated.

#### 2.5.2 Experimental design using Box-behnken

Three different levels variables and their interactions which influence maximal melanin production and cell biomass were analysed by response surface methodology, using Box-behnken design. 17 experimental trials constructed using two different coded levels (+1, -1) and its mean using temperature, pH and L-tyrosine as variables. All the experiments were done in triplicates and average cell biomass and melanin production were estimated as Reponses 1 and 2.

**Table 1:** Selected variables and their different levels.

| Factor | Name | Units | Minimum (+1) | Maximum (-1) | Mean |
| --- | --- | --- | --- | --- | --- |
| A | Temperature | <sup>0</sup>C | 30 | 35 | 32.50 |
| B | pH |  | 8 | 9 | 8.50 |
| C | L-tyrosine | g/L | 6 | 8 | 7.00 |

All the experiments were performed in triplicates for all the set of combinations generated from the design matrix with a total of 17 sets of experiments. The results were fitted on to a quadratic polynomial model by applying multiple regression analysis. The responses were predicted for given two levels (+1 and -1) of coded factors and its mean. The relative impact of factors was identified through comparison of factor coefficients.

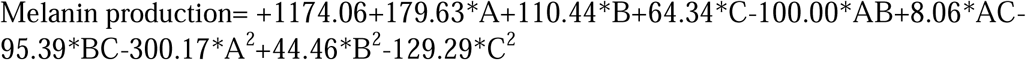

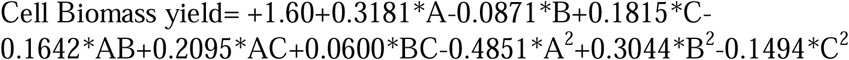

### 2.6 Microbial synthesis and characterization of melanin nano particle

The bacterial melanin obtained was sonicated at 30% amplitude, pulsing every 15 seconds for a total of 30 minutes in deionized water. The sonicated sample was then centrifuged at 18,000 rpm for 15 min to separate the mono sized nano melanin. The nano-melanin so obtained was characterized using TEM, particle size analyser and stability check was performed using Zeta potential (Mathew *et al*., 2022).

### 2.7 Anti-inflammatory property

The reaction mixture (0.5 ml) consisted of 0.4 ml bovine serum albumin (3% aqueous solution) and varying concentrations of test sample. The samples were incubated at 37°C for 20 min and 2.5 ml phosphate buffered saline (pH 6.3) was added to each tube and then heated at 80°C for 10 min. The absorbance was measured using spectrophotometer at 660nm.The percentage inhibition of protein denaturation was calculated as follows:

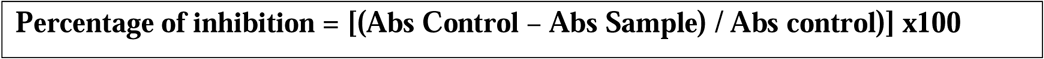

### 2.8 Wound healing potential

Wound healing potential of Nano sized melanin particles were performed in L6 cell line by scraped cell monolayer in a straight line by using 200 µL pipette tip. Incubated it in tissue culture incubator at 37 ^0^C for 48 h. The cell lines were visualised using phase contrast microscope till the closure of the wound.

## 3. Results and discussion

### 3.1 Biochemical characterization of bacterial strain

This marine aerobic Gram-negative rod-shaped bacteria BTCZ305 was characterized biochemically. It was Indole, catalase and oxidase positive and Methyl red, Voges-Proskauer and citrate negative. The utilization of different carbon sources including glucose, lactose, sorbitol, mannitol, Rhamnose and sucrose were performed. Of this sucrose responded positively.

### 3.2 Molecular identification through 16S DNA riboprofiling

16S RNA sequencing was carried out and the sequences were submitted in Genbank with Accession No. MW479156 Fig 1(A)(Mathew *et al*., 2022)

**Fig 1:**
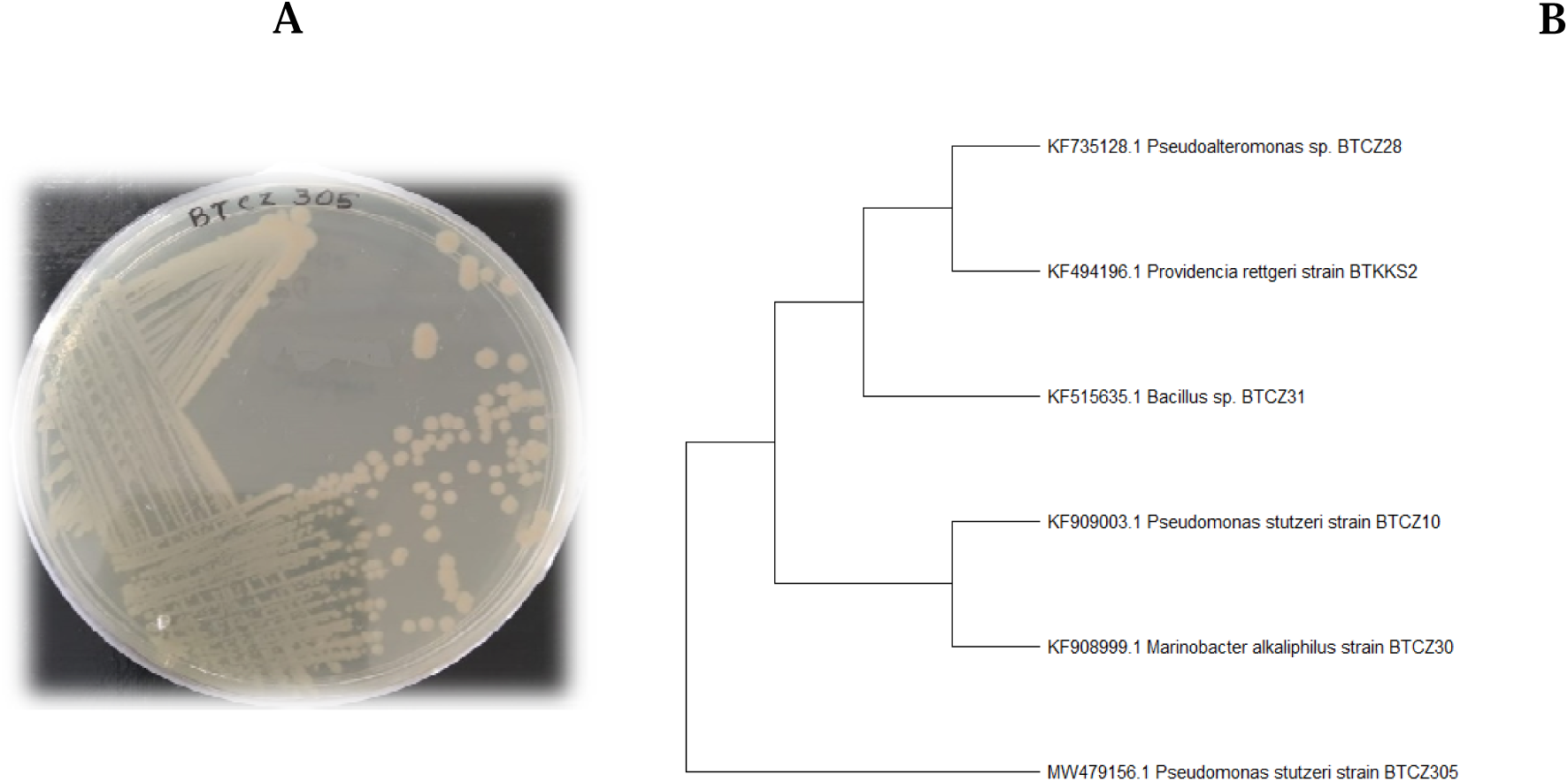
(A) Nutrient agar plate showing *Pseudomonas stutzeri* BTCZ 305. (B) Evolutionary relationships of taxa were demonstrated as phylogenetic tree using Neighbor-Joining method in Mega X software.

### 3.3 Time course study on melanin production

The time course study was conducted up to 216 h at a regular interval of 12 h in YMM medium and maximum melanin production was achieved at the stationary phase of bacterial growth. Melanin production built up slowly during exponential phase and attained maxima at the onset of stationary phase (Fig 2).

**Fig 2:**
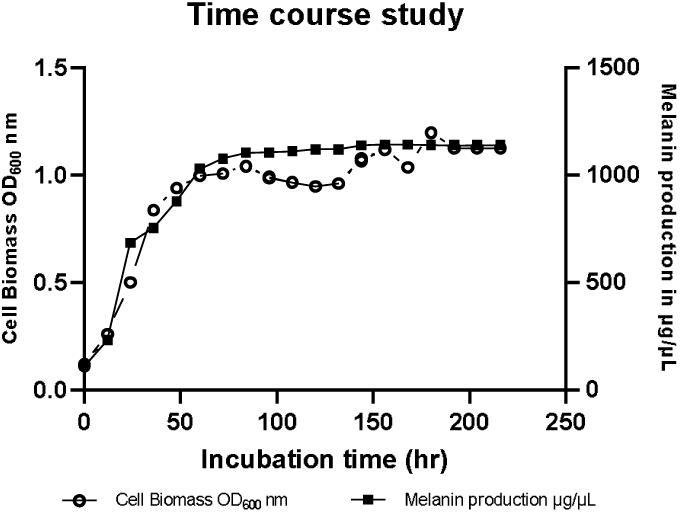
Time course study on cell biomass concentration and melanin production by *Pseudomonas stutzeri* BTCZ 305.

### 3.4 Formulation of fermentation media through OFAT method using most influencing physicochemical factors

#### 3.4.1 Effect of pH

Maximum melanin production and cell biomass concentration were observed in alkaline pH range (7-9) and diminishing towards acidic range. One study conducted by Aghajanyan *et al*., 2005 suggested that pH range from 7-8 was required for *Bacillus thuringiensis* melanin synthesis. Melanin from marine Streptomyces sp. (MVCS13) showed (Fig 3(A)) maximum melanin production at a pH of 7.4 (Sivaperumal *et al*.,2014). Some previous research show evidence of bacterial melanin stable only in the pH range of 4.0-11.0 (Wang *et al*.,2006). Among different parameters studied pH of the growth medium plays a crucial role in the stability of melanin granule in the medium. pH of the medium helps in aggregation as well as disintegration of the granules.

**Fig 3:**
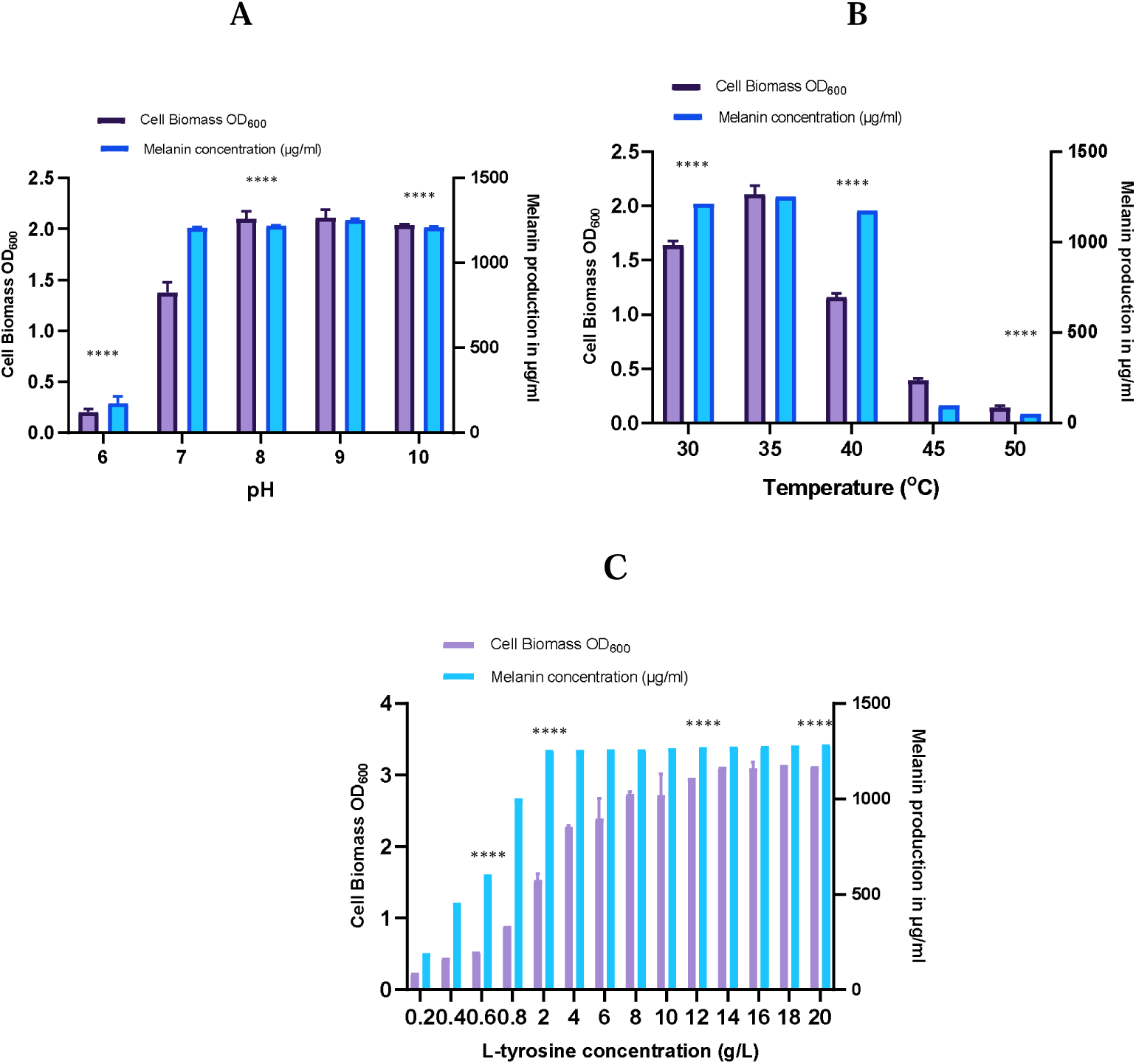
Effect of (A) pH (B) incubation temperature and (C) L-tyrosine concentration on melanin production by *Pseudomonas stutzeri* BTCZ305. Statistically analysed using Graph pad prism 8 in Windows 10. The results are means of three replicates +/-SD. It was found that P<0.0001(****) and all the results were statistically significant.

#### 3.4.2 Effect of temperature

Incubation temperature is also a critical factor in the growth of bacteria. Experiments in triplicates were performed on various temperature ranging from 30-50 °C. Results indicate that maximum melanin production and cell biomass was noted at 35 °C9Fig 3(B)). As the incubation temperature was further increased decrease in melanin production was also observed. In a similar study conducted on *Bacillus thuringiensis* (Ruan *et al*., 2004) the high melanin yield was obtained when temperature elevated to 42 ^0^C. In a recent study conducted on actinobacteria *Dietzia schimae* (Eskandari *et al*., 2020) maximum amount of melanin was obtained at an optimum temperature of 32 ^0^C.

#### 3.4.3 Effect of L-tyrosine concentration

L-tyrosine act as the sole source of carbon and Nitrogen which influence melanin production and cell biomass concentration. Concentration ranging from 0.2-20 g/L were studied to determine the optimum concentration (Fig 3(C)). Results showed that increase in L-tyrosine increases melanin production which helps the cells to attain stationary phase at reduced time interval.

### 3.5 Statistical media optimization through Response surface methodology and Box-behnken DOE

Statistical media optimization using three different selected factors were performed using Box-behnken design. 17 trials were constructed using different combinations of all these factors and maximum melanin production and cell biomass was estimated (Inyang *et al*.,2020; Thite et al.,2020).

**Table 2:**
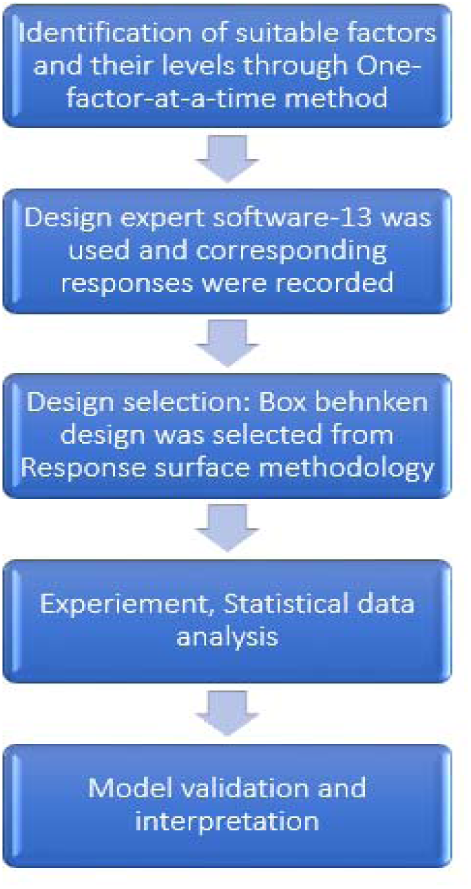
RSM method adopted for Cell biomass determination and Melanin production.

**Table 3:**
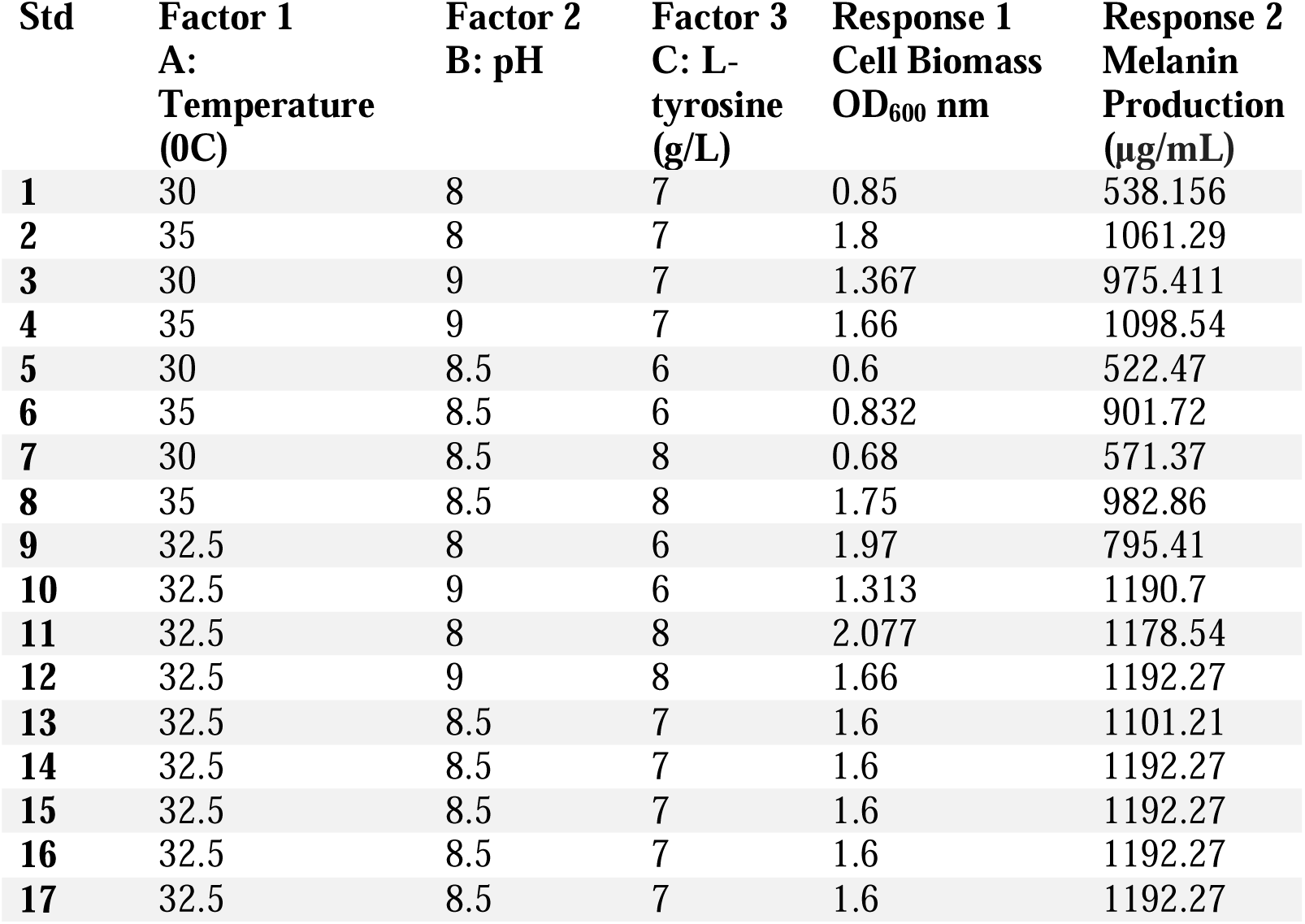
Box-behnken design showing Cell biomass and melanin production.

| Std | Factor 1<br>A:<br>Temperature<br>(0C) | Factor 2<br>B: pH | Factor 3<br>C: L-tyrosine<br>(g/L) | Response 1<br>Cell Biomass<br>OD<sub>600</sub> nm | Response 2<br>Melanin<br>Production<br>(µg/mL) |
| --- | --- | --- | --- | --- | --- |
| 1 | 30 | 8 | 7 | 0.85 | 538.156 |
| 2 | 35 | 8 | 7 | 1.8 | 1061.29 |
| 3 | 30 | 9 | 7 | 1.367 | 975.411 |
| 4 | 35 | 9 | 7 | 1.66 | 1098.54 |
| 5 | 30 | 8.5 | 6 | 0.6 | 522.47 |
| 6 | 35 | 8.5 | 6 | 0.832 | 901.72 |
| 7 | 30 | 8.5 | 8 | 0.68 | 571.37 |
| 8 | 35 | 8.5 | 8 | 1.75 | 982.86 |
| 9 | 32.5 | 8 | 6 | 1.97 | 795.41 |
| 10 | 32.5 | 9 | 6 | 1.313 | 1190.7 |
| 11 | 32.5 | 8 | 8 | 2.077 | 1178.54 |
| 12 | 32.5 | 9 | 8 | 1.66 | 1192.27 |
| 13 | 32.5 | 8.5 | 7 | 1.6 | 1101.21 |
| 14 | 32.5 | 8.5 | 7 | 1.6 | 1192.27 |
| 15 | 32.5 | 8.5 | 7 | 1.6 | 1192.27 |
| 16 | 32.5 | 8.5 | 7 | 1.6 | 1192.27 |
| 17 | 32.5 | 8.5 | 7 | 1.6 | 1192.27 |

The highest order polynomial where the additional terms are not significant and the model is not aliased were selected for further analysis. While considering both cell biomass and Melanin production a quadratic model maximising Adjusted R^2^ of 0.7825 for cell biomass and 0.5922 for melanin production were selected (Zeng *et al*.,2011; Ojha *et al*.,2020)

**Table 4:**
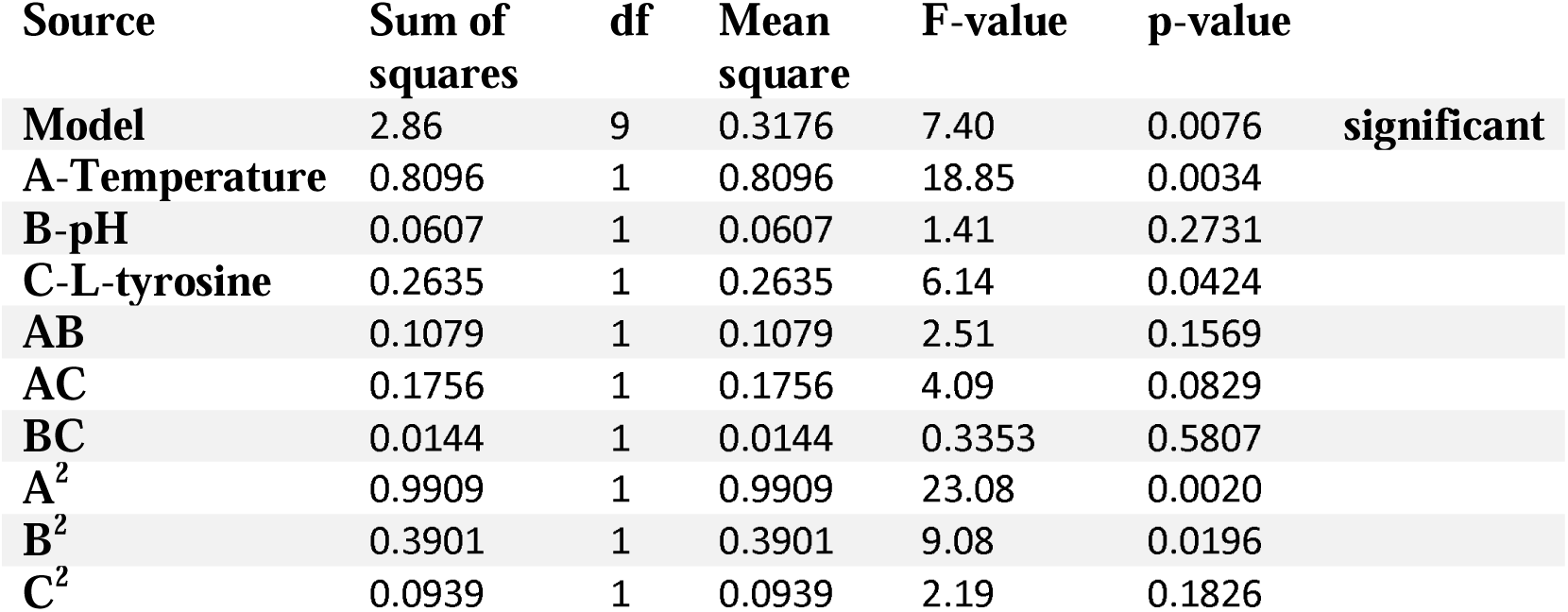
ANOVA table for cell biomass.

| Source | Sum of squares | df | Mean square | F-value | p-value |  |
| --- | --- | --- | --- | --- | --- | --- |
| **Model** | 2.86 | 9 | 0.3176 | 7.40 | 0.0076 | **significant** |
| **A-Temperature** | 0.8096 | 1 | 0.8096 | 18.85 | 0.0034 |  |
| **B-pH** | 0.0607 | 1 | 0.0607 | 1.41 | 0.2731 |  |
| **C-L-tyrosine** | 0.2635 | 1 | 0.2635 | 6.14 | 0.0424 |  |
| **AB** | 0.1079 | 1 | 0.1079 | 2.51 | 0.1569 |  |
| **AC** | 0.1756 | 1 | 0.1756 | 4.09 | 0.0829 |  |
| **BC** | 0.0144 | 1 | 0.0144 | 0.3353 | 0.5807 |  |
| **A<sup>2</sup>** | 0.9909 | 1 | 0.9909 | 23.08 | 0.0020 |  |
| **B<sup>2</sup>** | 0.3901 | 1 | 0.3901 | 9.08 | 0.0196 |  |
| **C<sup>2</sup>** | 0.0939 | 1 | 0.0939 | 2.19 | 0.1826 |  |

Here in Analysis of variance for cell biomass, the Model F-value of 7.40 implies that the model is significant. There is only a 0.76% chance that an F-value this large could occur due to noise. P-value less than 0.0500 indicate that the model terms are significant. The model with an insignificant lack of fit was selected.

**Table 5:**
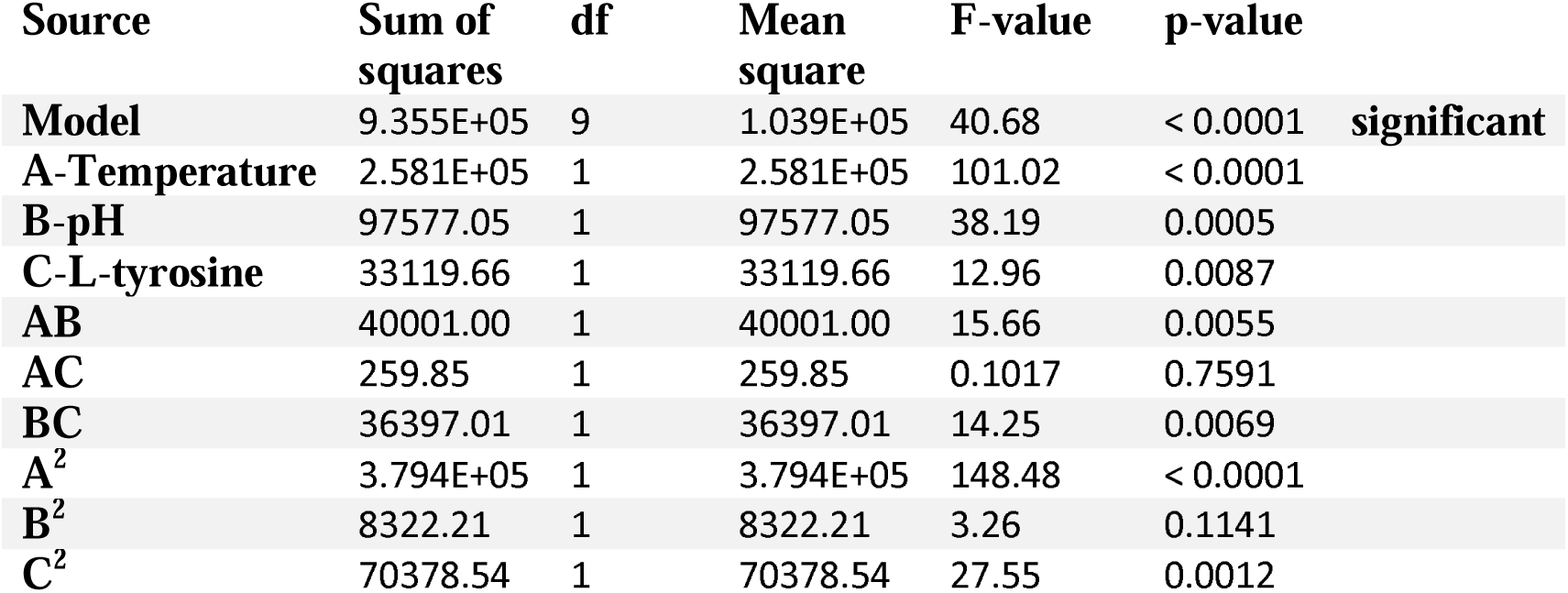
ANOVA table for Melanin production.

In the case of Analysis of variance for melanin production, the Model F-value of 40.68 implies the model is significant. There is only a 0.01% chance that an F-value this large could occur due to noise. P-value less than 0.0500 indicate model terms are significant.

The lack of fit F-value of melanin production 2.26 implies that the lack of fit is not significant relative to the pure error. There is a 22.34% chance that the lack of fit F-value this large could occur due to noise. For the model to fit, non-significant lack of fit is preferred.

#### 3.5.1 Statistical analysis and model validation

The R^2^ values were 0.9048 for cell biomass and 0.9812 for melanin production, suggesting that the model could explain 90.48 and 98.12% of total variation in cell biomass and melanin production respectively.3 D surface and contour graphs were plotted representing the interactive effect of different factors which leads to maximum melanin production and cell biomass concentration.

**Fig 4.**
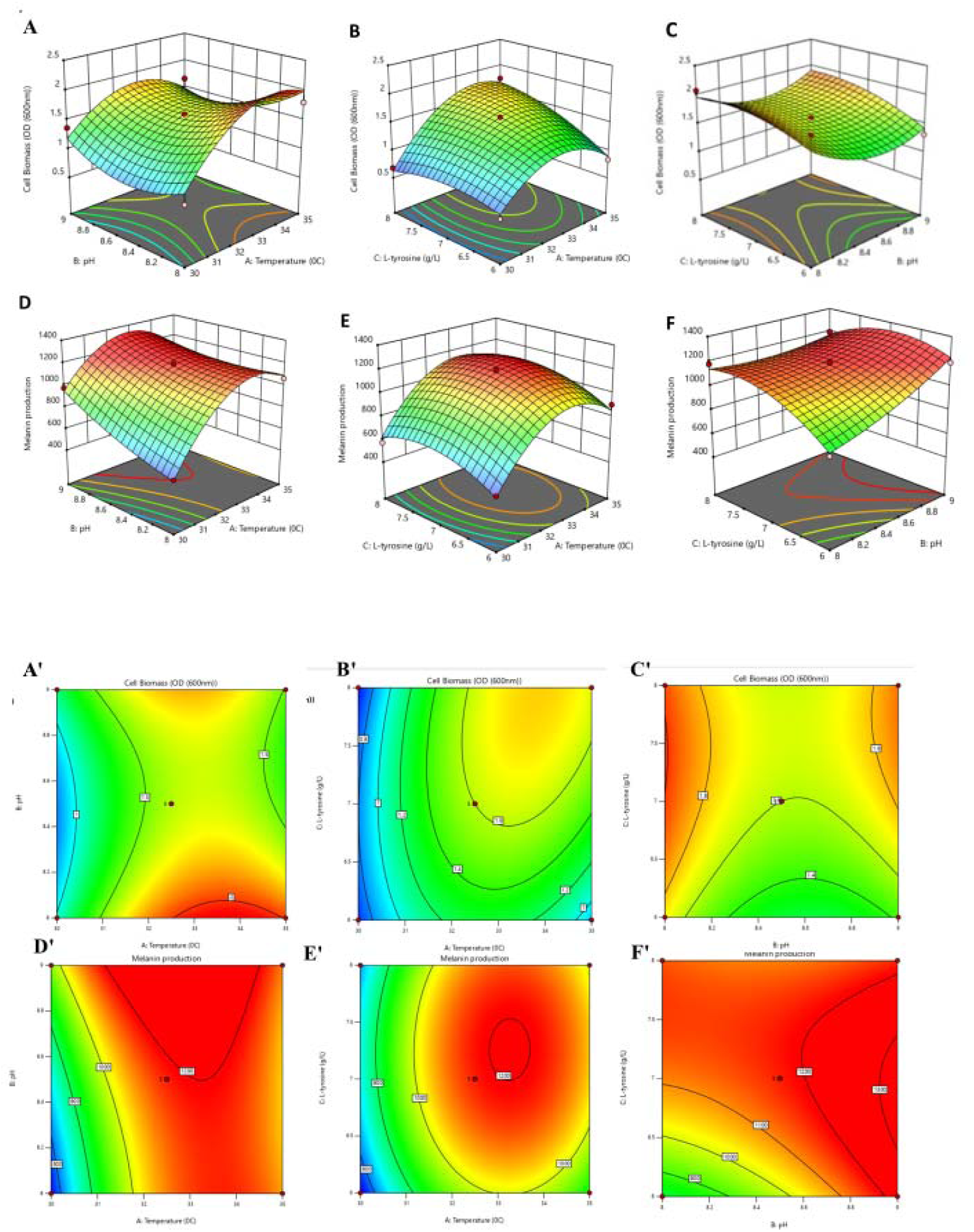
3D surface and contour plots representing the interactive effect of variables for both cell biomass and melanin production(A and A’) pH and temperature, (B and B’) L-tyrosine and temperature, (C and C’) L-tyrosine and pH, (D and D’) pH and temperature, (E and E’) L-tyrosine and temperature (F and F’) L-tyrosine and pH for *Pseudomonas stutzeri* strain BTCZ 305 were depicted.

#### 3.5.2 Experimental data validation

Three different variables differentially influence cell biomass yield and melanin production. Observed vs predicted results for cell biomass and melanin production were shown in fig 5(A)(B). gives confirmation on this assumption. The maximal cell biomass and melanin production was observed at a temperature of 32.5 ^0^C, pH of 8.5 and L-tyrosine concentration of 7 g/L.

**Fig 5.**
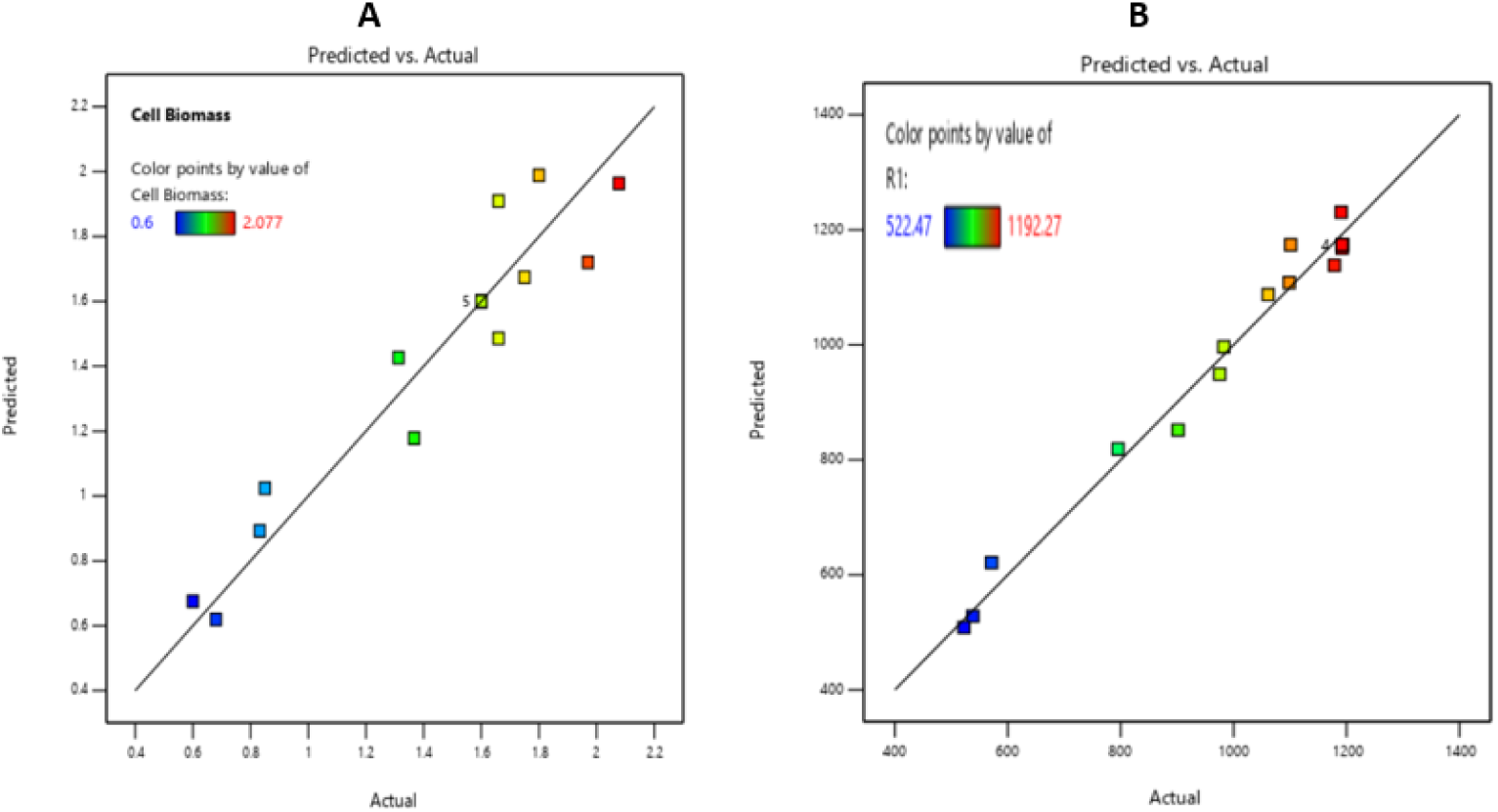
The actual vs predicted values were shown under optimised conditions of (A) Cell biomass and (B) Melanin production of *Pseudomonas Stutzeri* strain BTCZ 305

The optimized predicted values for maximal cell biomass and melanin production are 1.6 (OD_600_ nm) and 1174.06 μg/mL respectively at optimum levels of temperature 32.5 ^0^C, pH 8.5 and L-tyrosine concentration 7 g/L. The actual values obtained for cell biomass yield and melanin production were 1.6 at OD_600_ nm and 1192.27 μg/mL respectively and are appreciably close to predicted values, thus validating the model. The data obtained was further validated through triplicated conformation experiments. An increase of 7.39-fold was observed in melanin production compared to unoptimized medium which authenticates the efficiency of Response surface methodology Box-behnken design.

### 3.6 Microbial synthesis and characterization of melanin nano particle

The melanin obtained after centrifugation and acid precipitation was sonicated and evaluated using diffractive light scattering, zeta potential and Transmission electron Microscopy to determine particle size distribution, stability and size range respectively of these nano particles.

#### 3.6.1 UV-visible spectroscopy

UV-Visible absorbance spectrum of melanin was generated with an absorbance range of 200-800 nm. The absorbance maximum was observed between 200-250nm, but diminishes towards the visible region and it was due to the actual complex structure of melanin (Tarangini *et al*., 2014). The absorbance peak obtained from BTCZ305 showed maximum absorbance at 211 nm(Fig 6(A)). This result shows similarity towards the melanin obtained from purified *Chroogomphus rutilus* melanin which had maximum absorption peak at 212 nm (Hu *et al*., 2015). The absorption peak obtained through UV-Vis spectrometer also varies among melanin synthesised from different species.

**Fig 6.**
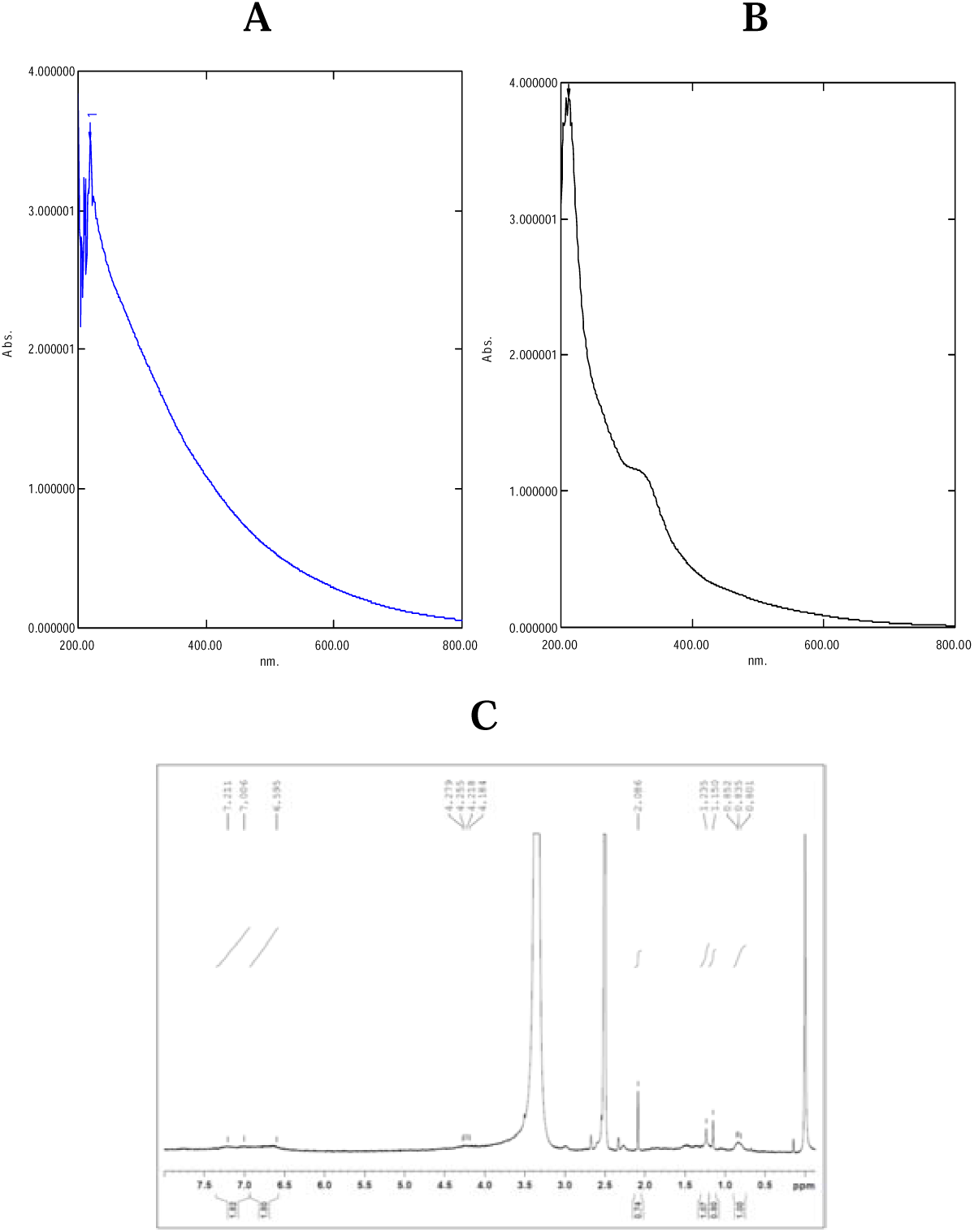
UV-visible absorption spectrum (200-800 nm) of the purified melanin pigment of (A)DOPA Melanin; (B) BTCZ305 (C) ^1^H NMR

**Fig 6.**
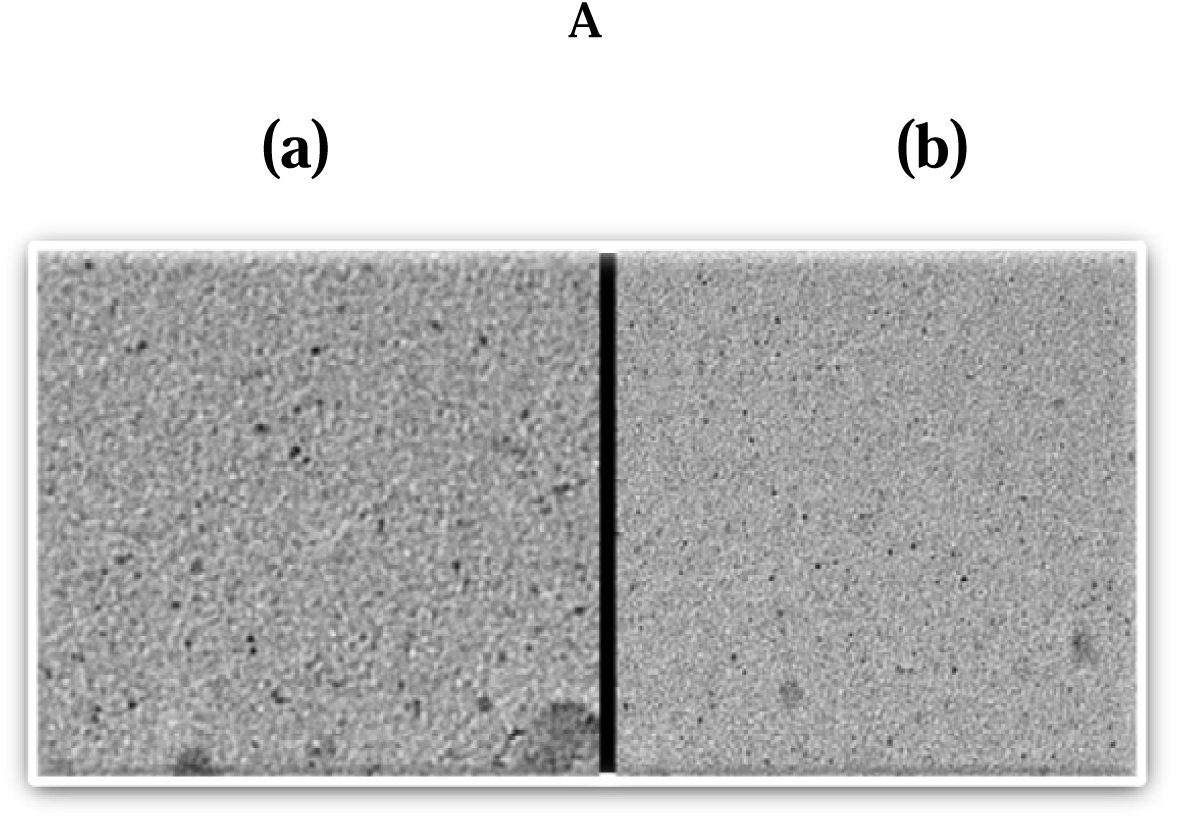
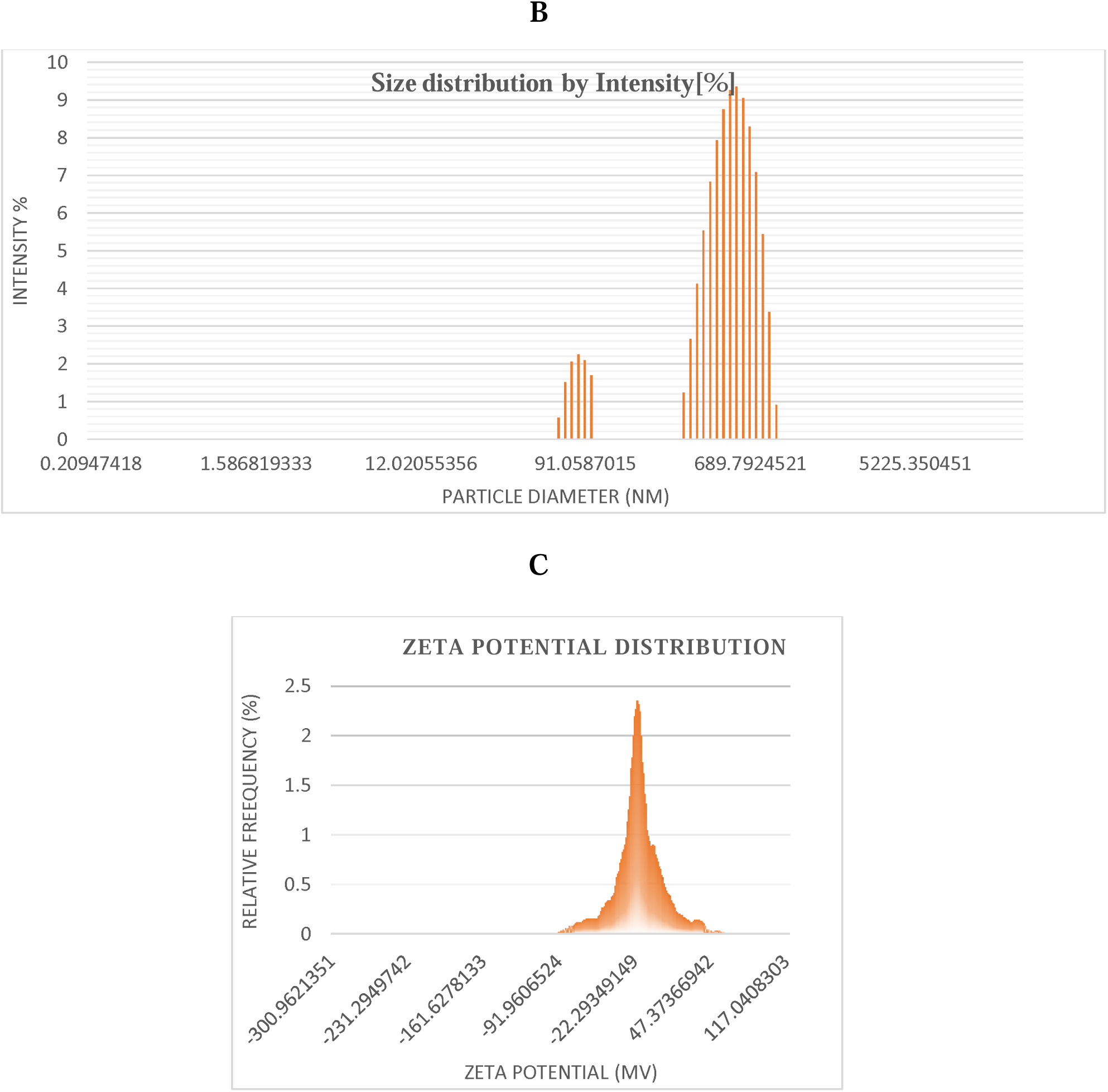
Characterization of melanin nano particle using (A) TEM (B) Particle size analysis (C) zeta potential

#### 3.6.2 ^1^H NMR

NMR spectrum showed (Fig 6(C) resonance between 0.8 and 4.2 ppm. The resonance between 8.0 and 6.0 ppm are attributed to aromatic functionality, between 3.0 and 4.3 ppm are assigned to protons attached to N or O. Resonance 6.595, 7.006 and 7.211ppm for BT CZ 305 are assigned to indole and pyrrole repeat units of the melanin polymer (Katritzk *et al*., 2002).

#### 3.6.3 TEM

Transmission electron microscope showing ∼5-7 nm of mono sized nano melanin particle with spherical geometry were shown at different magnifications. Nano melanin granules under different magnifications (50 and 100 nm) were shown below. [Fig 6. A(a)(b)].

#### 3.6.4 Particle size analysis and Zeta potential distribution

Dynamic light scattering (DLS), often referred to as photon correlation spectroscopy (PCS), is a common technique for determining particle size in colloidal suspensions (Hoo *et al*.,2008) depending on the Brownian motion of a particle. It was inferred from the result that the-melanin granules synthesised by *Pseudomonas stutzeri* strain BTCZ305 exhibited nano sized distributions in the range of 98.74 nm (with a smaller intensity of 2.24% and width of 32.89) and 953.72 nm (with a smaller intensity of 9.3 % and width of 646.85) with a mean hydrodynamic diameter of 557.7 nm. The increase in diameter was due to the existence of water molecules making non-covalent interaction with the melanin nano granules. The zeta potential was -22.3 +/-1.5 mV (Fig 6 (b)). The negative zeta potential value indicated the electrostatic stability of this molecule in colloidal suspension. The increased stability with nano-sized distribution makes it an effective candidate in several therapeutic applications.

### 3.7 Anti-inflammatory property

The anti-inflammatory property of melanin at its different concentrations were depicted in fig. From the results it was observed that increase in concentrations of melanin increases % inhibition of protein denaturation which proves the anti-inflammatory potential of this biomolecule (Mizushima *et al*.,1968; Sakat *et al*.,2010).

### 3.8 Wound healing potential

The Melanin nano particle synthesised by *Pseudomonas stutzeri* BTCZ 305 has wound healing property. In the control experiment even after 36 hours of incubation, the wound remained more or less similar. While in treated sample the wound was fully healed at 36 hours. Also, the efficiency and speed of wound healing were also affected by the concentration of the compound (Liang *et al*., 2007).

**Fig 7.**
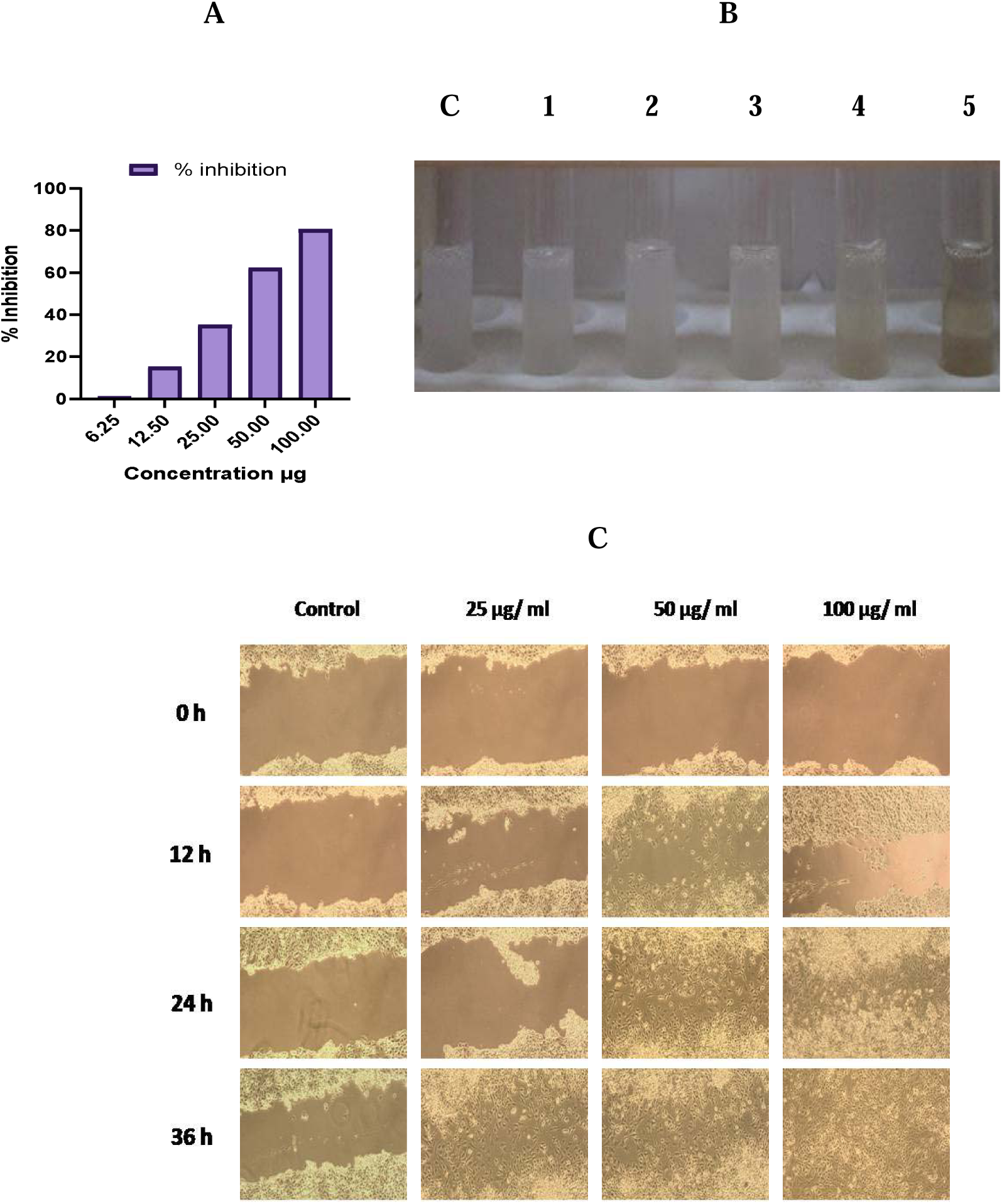
(A)(B) Anti-inflammatory property of Melanin was demonstrated as % inhibition of protein denaturation. (C) The wound healing potential of melanin on L6 Cell line at different time interval was compared.

## 4 Conclusion

Cultural condition favouring maximum cell biomass and melanin production were optimized through one-factor-at-a-time method and statistical method like Response surface methodology was employed for determining the most influencing factors. 17 different combinations of all the three factors were performed using Box-behnken design. A quadratic regression model was built with R^2^ values of 0.9048 for cell biomass and 0.9812 for melanin production and maximum melanin production was estimated to be 1192.27 μg/mL. Cell biomass and melanin production obtained experimentally coincides with the predicted value and hence the model was validated. The nano structural distribution of the molecule, its stability and size were determined and the applicability of the molecule in anti-inflammation and wound healing was also evaluated.

## 5. Acknowledgements

The first author acknowledge the support of Council of Scientific and Industrial Research (CSIR), Govt. of India for granting fellowships during the study (CSIR Award no: 09/239(0535)2018-EMR-1).

## 7 Author contribution

The first author did all the experimental work and wrote the draft of the manuscript under the supervision of the corresponding author, who is her doctoral supervisor, and mentor, who modified the manuscript for publication.

